# Cortical microcircuit determination through global perturbation and sparse sampling in grid cells

**DOI:** 10.1101/019224

**Authors:** John Widloski, Ila R. Fiete

## Abstract

Under modern interrogation, famously well-studied neural circuits such as that for orientation tuning in V1 are steadily giving up their secrets, but quite basic questions about connectivity and dynamics, including whether most computation is done by lateral processing or by selective feedforward summation, remain unresolved. We show here that grid cells offer a particularly rich opportunity for dissecting the mechanistic underpinnings of a cortical circuit, through a strategy based on global circuit perturbation combined with sparse neural recordings. The strategy is based on the theoretical insight that small perturbations of circuit activity will result in characteristic quantal shifts in the spatial tuning relationships between grid cells, which should be observable from multi- single unit recordings of a small subsample of the population. The predicted shifts differ qualitatively across candidate recurrent network mechanisms, and also distinguish between recurrent versus feedforward mechanisms. More generally, the proposed strategy demonstrates how sparse neural recordings coupled with global perturbation in the grid cell system can reveal much more about circuit mechanism as it relates to function than can full knowledge of network activity or of the synaptic connectivity matrix.

## Introduction

Questions about the origin of the beautiful tuning curves often seen in sensory and cortical circuits have long consumed systems neuroscientists, both theorists who propose possible mechanisms, and experimentalists who search for them (Hubel and Wiesel, 1959). Indeed, the mechanisms underlying direction tuning in the retina and cortex and orientation tuning in V1 remain unresolved and closely studied (Rivlin-Etzion et al., 2012; Kim et al., 2014; Takemura et al., 2013; Ferster and Miller, 2000; Sompolinsky and Shapley, 1997). Basic questions, like whether orientation tuning is largely attributable to selective feedforward summation or lateral interactions, are not yet settled.

Here we propose that the grid cell system provides a unique opportunity for understanding the underpinnings of computation in cortical circuits. The unusual responses of grid cells present a challenge and simultaneously, an opportunity. The challenge is to understand how such a complex cognitive response is generated; the opportunity is the availability of versatile experimental tools and a rich set of relatively detailed models (Fuhs and Touretzky, 2006; Guanella et al., 2007; Burak and Fiete, 2009; Burgess et al., 2007; Hasselmo et al., 2007; Welday et al., 2011; Mhatre et al., 2012; Bush and Burgess, 2014; Hasselmo and Brandon, 2012; Navratilova et al., 2012) that are well-constrained by the very complexity of the grid cell response, to help meet the challenge.

The recent application of quantitative analyses to electrophysiological data reveals that the population activity of grid cells (within individual modules) is localized around a continuous low-dimensional (2D) manifold (Yoon et al., 2013; Fyhn et al., 2007), a finding that lends support to early models predicated on the idea of low-dimensional pattern formation through strong lateral interactions (Fuhs and Touretzky, 2006; Burak and Fiete, 2006; McNaughton et al., 2006; Guanella et al., 2007; Burak and Fiete, 2009), as well as other models in which grid cells receive location-coded inputs and through structured feedforward connections (with the possible addition of some lateral connectivity) generate grid-patterned responses (Kropff and Treves, 2008; Mhatre et al., 2012; Welday et al., 2011; Bush and Burgess, 2014).

These models are architecturally and mechanistically distinct in important ways, both large and subtle: they differ in whether grid cells perform velocity-to-location integration, in whether pattern formation originates wholly or partly within grid cells, and in the structure of their recurrent circuitry. Some of the structural differences within recurrent models which seem subtle have qualitative ramifications for how the circuit could have developed. Despite their differences, the models are difficult to distinguish on the basis of existing multiple single-unit activity records, because all of them produce grid-patterned outputs and exhibit approximate 2D continuous attractor dynamics. Worse, as we discuss at the end, neither complete single neuron-resolution activity records nor complete single synapse-resolution weight matrices will be sufficient to distinguish between proposed mechanisms.

We show how it is nevertheless possible to gain surprisingly detailed information about the grid cell circuit from a feasible experimental strategy that depends on global circuit perturbation and sparse neural recording. In this context, global means circuit-wide not brain-wide. The proposed strategy can allow the experimenter to discriminate between various distinct candidate mechanisms that are currently undifferentiated by experiment.

## Results

### Experimentally undifferentiated grid cell models

Let us begin by considering 2D recurrent pattern forming models, in which grid cells are assumed to integrate velocity inputs and output location-coded responses. Such recurrent pattern forming models are of three main types. The first are *aperiodic* networks (Burak and Fiete, 2009; Widloski and Fiete, 2014), Figure 1A. In these models, activity in the cortical sheet (when neurons are appropriately rearranged – note that topography is not required in these models or in the proposed experiments) is grid-like and therefore periodic, but the connectivity between cells is highly localized and not periodic. In other words, connectivity does not reflect the periodicity in the activity. Taking a developmental or plasticity perspective, this network model is somewhat unusual in that strongly correlated neurons (those with the same activity phase, within or across activity bumps) are not connected as might be expected from associative learning. So if this network architecture holds in the brain, it would suggest that associative learning is curtailed once pattern formation occurs. From a functional viewpoint, aperiodic networks can require careful tuning of input at the network edges to accurately integrate their velocity inputs (Burak and Fiete (2009), but not so in Widloski and Fiete (2014)).

**Figure 1:**
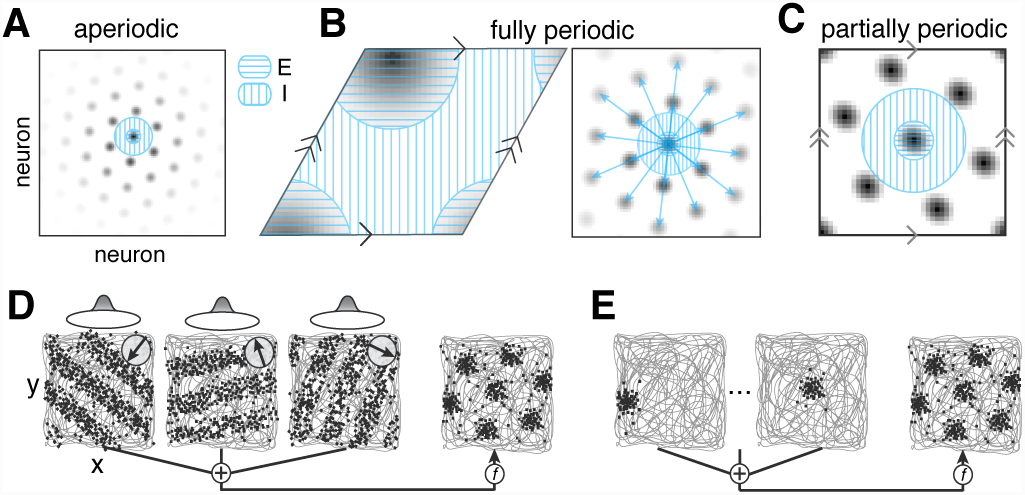
Mechanistically distinct models not distinguished by existing data. (A-C) Recurrent 2D pattern-forming models: Activity in the cortical sheet (gray; darker indicates more activity) and the outgoing recurrent weights from a single representative cell (blue green regions centered on the cell of origin). (A) Aperiodic network: Aperiodic boundary conditions and “local” connectivity that is not determined by activity phase so that connectivity does not reflect the periodicity in activity. (B) Fully periodic network: Connectivity period equals activity period, with periodic boundary conditions on a rhombus. The two networks shown (left: single-bump network; right: multi-bump network with all bumps identified by allowing for strong recurrent connections between cells of the same activity phase) are mathematically identical. We refer to both as a single-bump network. (C) Partially periodic network: “Local” connectivity (in same sense as in (A)), with opposite edges of the cortical sheet identified so that the network boundary conditions are periodic. (D-E) Feedforward and feedforward-recurrent models: Spatially tuned (post-path integration) inputs drive grid cells (gray: spatial trajectory; spikes: red). (D) The inputs, generated from ring attractor networks (ellipses above squares) that integrate different components of animal velocity indicated by the inset compasses, have stripe-like spatial tuning (as in Mhatre et al. (2012)). Feedforward summation followed by a nonlinearity produces grid-like responses (right). (E) Place-tuned inputs with selective feedforward summation, and in some models, lateral interactions, drive grid-like responses.

The second type are *fully periodic* networks, Figure 1B (Guanella et al., 2007; Burak and Fiete, 2006; Fuhs and Touretzky, 2006; Pastoll et al., 2013; Brecht et al., 2014; Widloski and Fiete, 2014). In these, network connectivity is itself periodic (the network has periodic boundary conditions on a rhombus), and the connectivity period equals the activity period: they each have a single period over the network (left, Figure 1B). An alternative version of the fully periodic network is to consider an aperiodic network with multiple activity bumps, but in which neurons at the centers of all the activity bumps are synaptically coupled. These two views of a fully periodic network are mathematically equivalent. Developmentally, the latter may be constructed from an aperiodic network (with multiple activity bumps) by application of associative learning post pattern-formation, so that neurons with similar phase but in different bumps end up recurrently coupled (right, Figure 1B).

The third type of the recurrent pattern forming networks is *partially periodic*, Figure 1C (Burak and Fiete, 2009). In these, as in the aperiodic networks, the bulk connectivity is local, so that connectivity does not reflect the periodicity of the population activity patterns. However, opposite edges of the cortical sheet are identified so that the network is effectively a torus. From a developmental perspective, these networks are the strangest: bulk connectivity does not reflect the periodic activity but the boundary condition requires knowledge of it (Figure S1).

Next come a variety of models in which grid cells are the result of feedforward summation of inputs that are already spatially tuned (Kropff and Treves, 2008; Welday et al., 2011; Mhatre et al., 2012; Bush and Burgess, 2014). Functionally, these models suggest that path integration occurs upstream of grid cells, in different low-dimensional attractor networks (Welday et al., 2011; Mhatre et al., 2012; Bush and Burgess, 2014). (In Kropff and Treves (2008), the origin of spatial tuning in the inputs is not directly modeled; if the assumed place-cell like inputs are based on path integration then the model will display low-dimensional dynamics, so we will consider the model under this assumption). Some of these models additionally include recurrent weights in the grid cell layer (Kropff and Treves, 2008; Mhatre et al., 2012). We will call all these *feedforward* models.

### A perturbation-based probe of circuit architecture

The conceptual idea for differentiating between recurrent models of grid cells depends on multi-single unit grid cell recording before and after a global perturbation of the network. The idea is as follows. If population activity patterning in the neural sheet is due to aperiodic recurrent connections, then globally increasing the gain of recurrent inhibition or the time-constant of neurons in network models is predicted to increase the period of stable patterns in the cortical sheet, Figure S2. These effects, not predicted by linear stability analysis, exist in simulation of dynamical models (Widloski and Fiete, 2014; Burak and Fiete, 2009) and can be analytically derived by considering nonlinear effects (Widloski and Marder, unpublished observations).

Following the global perturbation, two cells originally in adjacent peaks of the population activity pattern and thus at the same phase of the population pattern (Figure 2A, blue), no longer will be (Figure 2A-B, red). Call the shift in pattern phase between cells in neighboring peaks one quantum (Figure 2A, circle and square, red versus blue). Then the shift in pattern phase between cells previously of the same phase and separated by exactly *K* peaks will be *K* quanta (Figure 2A, circle and triangle, red versus blue; explicit phase plot in Figure 2B). Across all cell pairs in the population, the shifts in phase will be quantized and will reach a maximum value of *M* quanta (or a full phase cycle, whichever is smaller), where *M* is the number of bumps in the population pattern.

**Figure 2:**
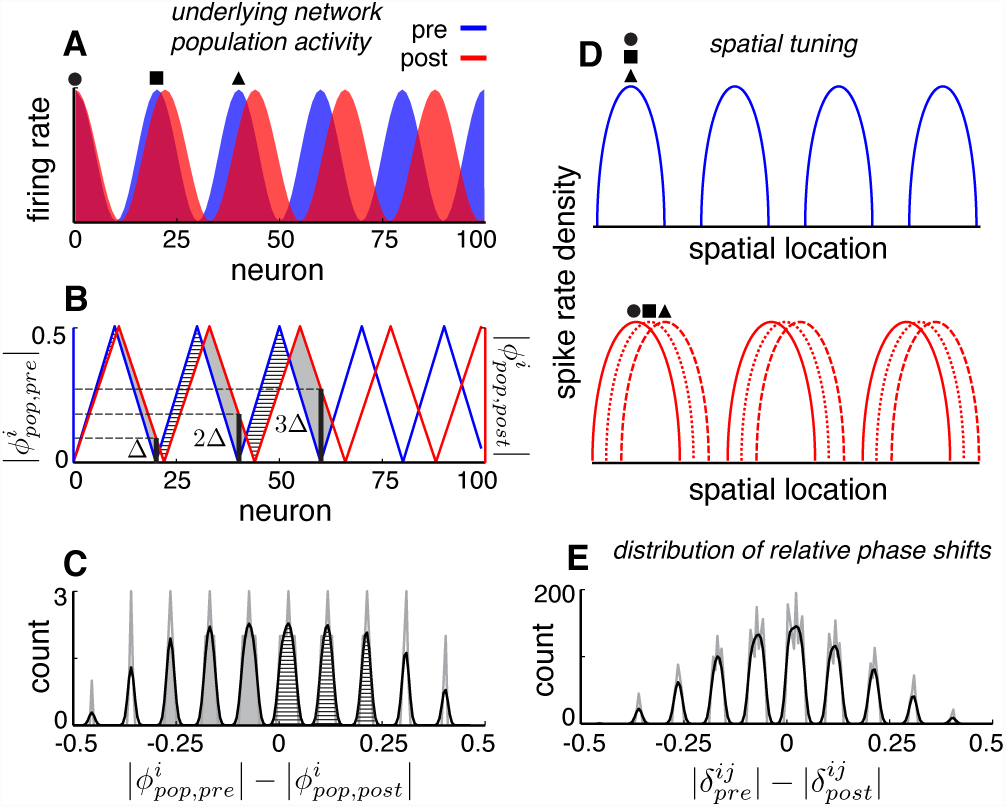
Global perturbation and phase shift analysis can reveal detailed features of population patterning. (A) Schematic (not a dynamical neural network simulation) of population activity in a 1D aperiodic grid cell network (blue) before perturbation and (red) after a 10% pattern expansion (*α* = 0.1; *λ*_*pop,pre*_ = 20 neurons). For illustration, cells are ordered topographically based on local connectivity and pattern expansion is centered at the left network edge. Circle, square, triangle: *pop* cells that shared the same phase no longer do post-expansion. (B) The pattern phase (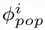 see Experimental Procedures) of cells (i) in the network, pre- (blue) and post- (red) perturbation. Cells exactly K peaks apart in the population pattern exhibit shifts in population phase equal to *K*Δ, where Δ is the quantal phase shift. (C) The histogram of shifts, pre- to post-perturbation, in the pattern phases of all cells (n=100). Gray line: raw histogram (200 bins). Black line: smoothed histogram (convolution with 2-bin Gaussian). Negative (positive) phase shifts are from gray-shaded (vertically-striped) areas in (B). (D-E) Shift distributions for pattern phase (experimentally inaccessible) carry over to shift distributions for relative spatial tuning phase (experimentally observable). (D) The circle, square, and triangle cells originally have identical spatial tuning (schematic in blue), but postperturbation are no longer co-active thanks to shifts in the population pattern (as in A) and thus also exhibit shifted spatial tuning curves (red). The shift for a pair is proportional to the number of activity bumps between them in the original population pattern. (E) Histogram of relative phase shifts (DRPS; gray). A relative phase shift (*δ*^*ij*^; see Experimental Procedures) between a pair of cells *i*, *j* equals the change in their relative spatial tuning phase post-perturbation. Black: smoothed version. There are *n* = (100 choose 2) samples because relative phase is computed pairwise.

Suppose the perturbation induces at most small phase shifts between all bumps of the population pattern (that is, 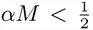, where 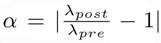 is the perturbation stretch factor, and *λ*_*pre*_, *λ*_*post*_ are the population pattern periods preand post-perturbation, respectively; see Figure S3). Then the number of peaks in the distribution of pattern phase shifts, Figure 2C, will equal twice the number of bumps in the underlying population pattern, Figure 2A. In other words, the number of peaks in the distribution of pattern phase shifts can specify the number of bumps in the population.

However, the construction of this distribution relies on experimentally difficult-to-access quantities, namely the population pattern phase for each cell. If the network were topographically organized, this would be relatively simple to extract from a snapshot of network activity. If the network is not topographically organized, it is possible to obtain estimates of phase similarity or phase distance magnitudes between cells from patterns of coactivation or correlation across snapshots of the population activity, but such a scalar activity similarity measure cannot yield 2D phase in a 2D network.

The utility of our proposed strategy arises because the distribution of shifts in the *population pattern phase* across cells is mirrored in the distribution of shifts in the *relative phase of spatial tuning* across cells (Figure 2D). We illustrate this point in 1D, but the same idea carries directly over to 2D (Figure S4). The relative spatial tuning phase is derived from the spatial tuning of simultaneously recorded cell pairs. Cell pairs with zero relative phase in their spatial tuning pre-perturbation (because they fell on the same phase of the population pattern, albeit in different bumps) will exhibit postperturbation shifts in relative phase that, like the shifts in the population phases, will be quantized, and for small changes in population period will be proportional to the number of bumps separating that cell pair, Figure 2C-E. This predicted *distribution of relative phase shifts* (DRPS, Figure 2E) between neurons from an aperiodic network is a property of patterning in an abstract space, independent of how neurons are actually arranged in the cortical sheet.

In 2D, relative phase is a vector, measured along the two principal axes of the spatial tuning grid. The total number of bumps in the population pattern can then be read out as equal to a quarter of the product of the number of peaks in the two relative phase shift distributions (Figure S4).

### Relating network parameters to experimental parameters

Changes in the strength of recurrent inhibition in our model can be mapped, in the biological system, into changes in the gain of inhibitory synaptic conductances. Experimentally, this perturbation may be induced by locally infusing allosteric modulators that increase inhibitory channel conductances (e.g. benzodiazipines (Rudolph and Möhler, 2004); personal communication with C. Barry). Changes in the time-constant of our model neurons can be mapped to changes in the EPSP time-constant in the biological system. Experimentally, the EPSP time-constant is sensitive to temperature through the Arrhenius effect and can be lengthened by cooling (Katz and Miledi, 1965; Thompson et al., 1985; Moser and Anderson, 1994). However, cooling affects several other single-neuron properties. To assess what to expect experimentally from a temperature perturbation and how to correctly include temperature effects in simpler neural models, we performed network simulations with cortical Hodgkin-Huxley neurons (Pospischil et al., 2008) while implementing documented temperature-dependent changes in all ionic and synaptic conductances (Experimental Procedures, SI, and Figure S5). The effect of cooling on conductance amplitudes is to shrink the population period in an aperiodic network, but its effect on conductance time-constants is to expand the period. The net effect of cooling is an expansion because temperature changes have larger effects on conductance time-constants (larger Q10 factors) than amplitudes (smaller Q10 factors) (Hodgkin et al., 1952; Thompson et al., 1985). We therefore conclude that changes in temperature are reasonable to associate with changes in the time-constant of simple neuron models. To summarize, two global experimental perturbations capable of inducing population activity period changes are neuromodulatory infusions that alter the gain of recurrent inhibition and cortical cooling (Moser and Anderson, 1994; Long and Fee, 2008).

### Discriminating amongst recurrent architectures

Dynamical simulations of grid cell models reveal that the effects of the global perturbation will differ across recurrent network architectures, with consequently different predictions for the DRPS. In an aperiodic network, incremental global perturbation results in incremental expansion of the population activity pattern (Figure 3A, red, and Figure S2). Thus, the DRPS envelope will gradually and linearly widen with perturbation strength, and the separation between peaks will gradually grow (Figure 3B-C, red and Figure S2).

**Figure 3:**
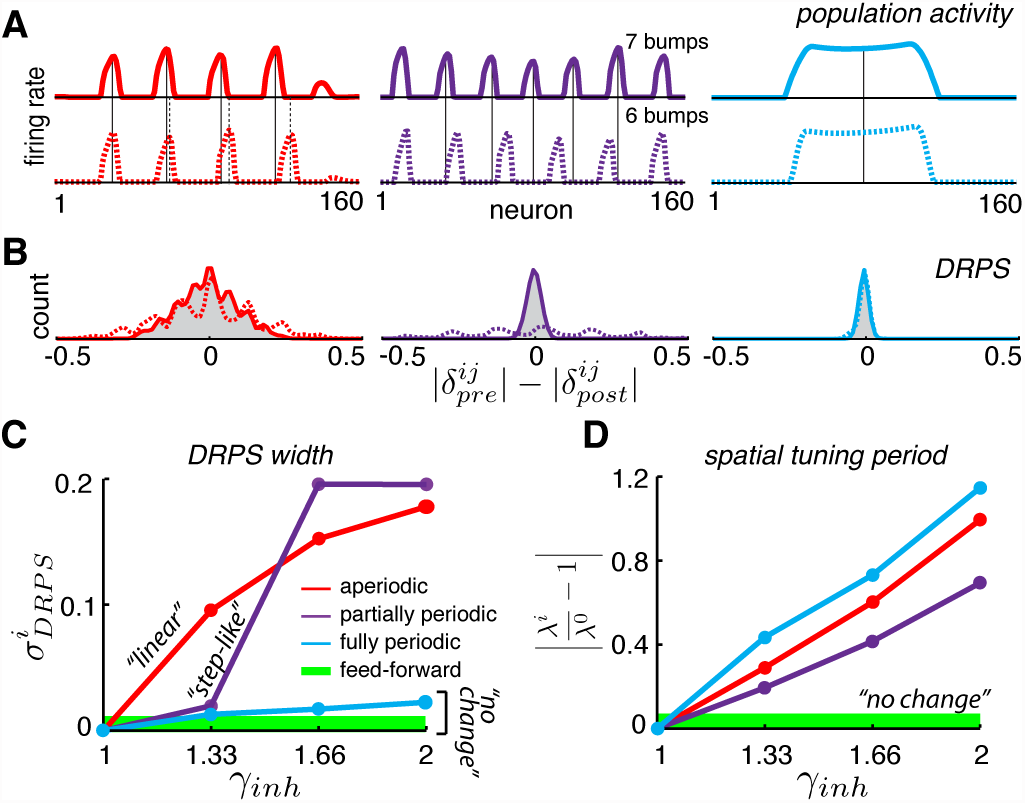
Effects of perturbation on recurrent and feedforward neural networks and predictions for experiment. (A-B) The effect of perturbing inhibitory weights in dynamical neural network simulations of aperiodic (column 1), partially periodic (column 2), and fully periodic (column 3) recurrent architectures (see Experimental Procedures for simulation details). (A) Population pattern preand post-perturbation (first and second rows, with *γ*_*inh*_ = 1 and 1.33, respectively). Vertical lines: bump centers in the unperturbed (solid) and perturbed (dotted) patterns. (B) DRPS relative to the unperturbed network. Solid line: perturbed network with *γ*_*inh*_ = 1.33; dashed line: larger perturbation of *γ*_*inh*_ = 1.66. (C) How the width of the DRPS, 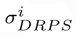, defined as the standard deviation of the DRPS, varies with perturbation strength. Thick green line: DRPS widths for feedforward networks (predicted, not from simulation). Note that, while the step-like shape of the DRPS width as a function of perturbation strength for the partially periodic network is general, the point at which the partially periodic network steps up will vary from trial to trial. (D) How the spatial tuning periods 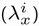 vary with perturbation strength in the different simulated recurrent networks and in a feedforward network (thick green line; predicted, not from simulation) (see SI for definition of spatial tuning period).

In a partially periodic network (with aperiodic local connectivity but with opposite boundaries connected), the number of bumps in the population activity pattern is constrained to be an integer. Thus, incrementally increasing the perturbation strength should result first in no change to the population activity period, and then a sudden change when the network can accommodate an additional bump (or an additional row of bumps in 2D, assuming the pattern does not rotate as a result of the perturbation; see Discussion) (Figure 3A, purple). Thus, incremental changes in perturbation strength should result in a stepwise change in population period and in the width of the DRPS envelope (Figure 3B-C, purple). Because the number of bumps has increased by a discrete amount, as soon as the DRPS changes, it will become maximally wide.

The fine structure of the DRPS will still be multimodal. However, counting peaks to estimate the number of bumps in the underlying population pattern will result in serious underestimation: when the pattern change is not incremental, there can be large changes in phase that are then lost in the DRPS, which is cut off at the maximal phase norm of 0.5 (Figure S3 and e.g. Figure 3B, compare peaks in the solid and dashed lines for small and large perturbations, respectively).

In the fully periodic network (Figure 1C), the same global perturbations that alter the population pattern period in the other recurrent networks (Figure 1A-B) are ineffective in inducing a corresponding change (Figure 3A, blue). This is because the periodic connectivity completely fixes the period of the pattern. Thus, the global perturbation will not affect the relative phase relationships between cells, and the DRPS is predicted to remain narrow, unimodal, and peaked at zero (Figure 3B-C, blue).

### Discriminating feedforward from recurrent architectures

If the spatial tuning pattern or pattern components are generated upstream of grid cells and inherited or combined by them through feedforward summation (Mhatre et al., 2012; Welday et al., 2011; Bush and Burgess, 2014), then perturbing the recurrent weights or the biophysical time-constant within only the grid cell layer is predicted to leave unchanged the population activity period, preserving the spatial tuning shapes and cell-cell relationships. As a result, the DRPS should be narrow and centered at zero, as in the case of a recurrent network with fully periodic connectivity (Figure 3C, green line).

In all recurrent model networks (Figure 1A-C), the *spatial tuning period* of cells is predicted to expand with the global perturbation, which induces a change in the efficacy with which feedforward velocity inputs shift the pattern phase over time (Figure 3D and Figure S6). This expansion in spatial tuning period with global perturbation strength is predicted to hold for all three recurrent network classes, and can be used as an assay of the effectiveness of the experimental manipulation, especially when there is no shift in the DRPS.

By contrast, in feedforward models integration occurs upstream of the grid cells and thus the spatial tuning period should remain unchanged with global perturbation (Figure 3D, green line). Response amplitudes should nevertheless change in the feedforward models, thus revealing whether the attempted global perturbations are in effect.

### Experimental feasibility of proposed method

We consider two key data limitations. First, it is not yet experimentally feasible to record from all cells in a grid module. Even a 100 cell sample would constitute a 1-10 % subsampling of the estimated module size. With present estimates that *<* 20 % of cells in a local patch in MEC are grid cells (Tang et al., 2014), the yield would be a meager 20 grid cells. Is this sufficient to observe the predicted quantal structure in a phase shift distribution, if it were present? Fortunately, the proposed method is tolerant to severe sub-sampling of the population: a tiny random fraction of the population (10/1600 cells) can capture the essential structure of the full DRPS, Figure 4A.

**Figure 4:**
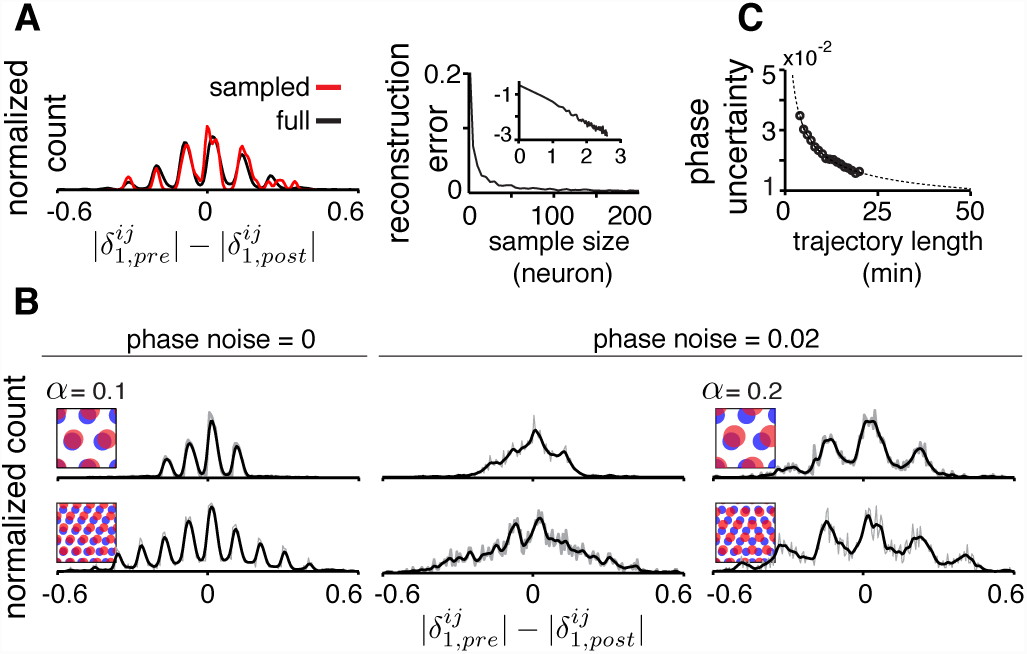
Measurement limitations and the resolvability of predictions under such constraints. (A) Left: The quantal structure of the DRPS is apparent even in small samples of the population (black: full population DRPS; red: DRPS computed from n=10 cells; both curves smoothed with 2-bin Gaussian; bins=200). Plotted is DRPS along first principal axis of the 2D phase (see Figure S4). Right: The L2-norm difference between the full and sampled DRPS as a function of number of sampled cells. Inset: log-log scale. (B) First and second columns: DRPS (200 bins; gray line: raw; black line: smoothed with 2-bin Gaussian) for different numbers of population pattern bumps along the first principal axis of the pattern and for different amounts of phase noise (noise is sampled i.i.d. from a gaussian distribution, 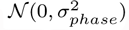, and added to each component of the relative phase vector, 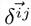; “phase noise” is the same as *σ*_*phase*_). Third column: Same as the second column, except for a larger stretch factor, *α* = 0.2. Note that the peak-to-peak separation has increased so that the individual peaks are discernible. However, for the 5 bump network in the second row, inferring the number of bumps in the underlying population pattern would lead to an underestimate, since *M × α* = 5 × 0.2 *>* 1*/*2. (C) The uncertainty (standard deviation) in estimating relative phase, for different amounts of data (data from Hafting et al. (2005)), from bootstrap samples of the full dataset (see SI for details). As expected, the decrease in uncertainty follows 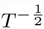 (gray). *Parameters*: *λ*_*pop,pre*_ = 40/3 neurons (A), = 20 neurons (B, top row), = 8 neurons (B, bottom row); α = 0.1;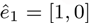; 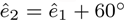; network size: 40 *×* 40 neurons.

Second, spatial tuning and relative phase parameters are estimated from neural responses during a random, finite exploration trajectory in which cells respond variably. Hence, spatial tuning parameters, including phase and relative phase, are only known with a degree of uncertainty. In tests that depend only on the width of the DRPS (e.g. Figure 3), this phase uncertainty is not a serious limitation.

However, more detailed questions about the number of bumps in the population pattern in an aperiodic network depend on estimating the number of DRPS peaks, and here phase estimation uncertainty can be problematic: phase uncertainty will merge together peaks in the DRPS, Figure S7. At very small perturbation strengths, the DRPS peak spacing (in the aperiodic network) increases with the stretch factor. Thus, the larger the perturbation, the more distinguishable the peaks at a fixed phase error, Figure 4B and Figure S7. Yet increasing the stretch factor is not without a tradeoff: The two-for-one relationship between number of peaks in the DRPS and the number of bumps in the population pattern per linear dimension holds when the total induced shift in phase is small for all bumps (as before, when 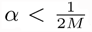, with *M* now equal to the larger of the number of bumps along the two principal axes of the population pattern), Figure 4B. At larger stretch factors, the number of peaks in the DRPS is smaller than twice the number of bumps along the corresponding dimension of the pattern, and the discrepancy can be substantial.

Fortunately, the DRPS is computed from the relative phases between cells, which remain stable in a fixed network (Yoon et al., 2013) (here fixed refers to the network while a given perturbation strength is stably maintained). This stability makes it possible to gain progressively better estimates of relative phase over time even if there is substantial drift in the spatial responses of cells, by computing the relative spatial phase over short snapshots of the trajectory then averaging together the relative phase estimates from different snapshots across a progressively longer trajectory (similar to the methods used in Yoon et al. (2013) and Bonnevie et al. (2013)).

To distinguish *M* = 5 bumps per linear dimension based on structure within the DRPS would require a stretch factor of no greater than *α* = 1*/*(2*M*) = 0.1, and phase noise must be reduced to at least 0.02 (Figure S7). Distinguishing 7 bumps would require *α ≤* 0.07 and a phase noise of smaller than about 0.01. Based on grid cell and trajectory data (accessed through <u>http://www.ntnu.edu/kavli/research/grid</u>cell-data), this would require an approximately 10 (50) -minute recording (Figure 4C).

The proposed method therefore has high tolerance to subsampling and a more limited tolerance to phase uncertainty. It will require longer-than-usual but still realistic amounts of spatial trajectory data with neural recordings to obtain adequately small error in relative phase estimation to test predictions that differentiate between models.

### A decision tree for experimental design

We lay out a decision tree with an experimental workflow for discriminating between disparate networks, all of which exhibit 2D continuous attractor dynamics (Figure 5).

**Figure 5:**
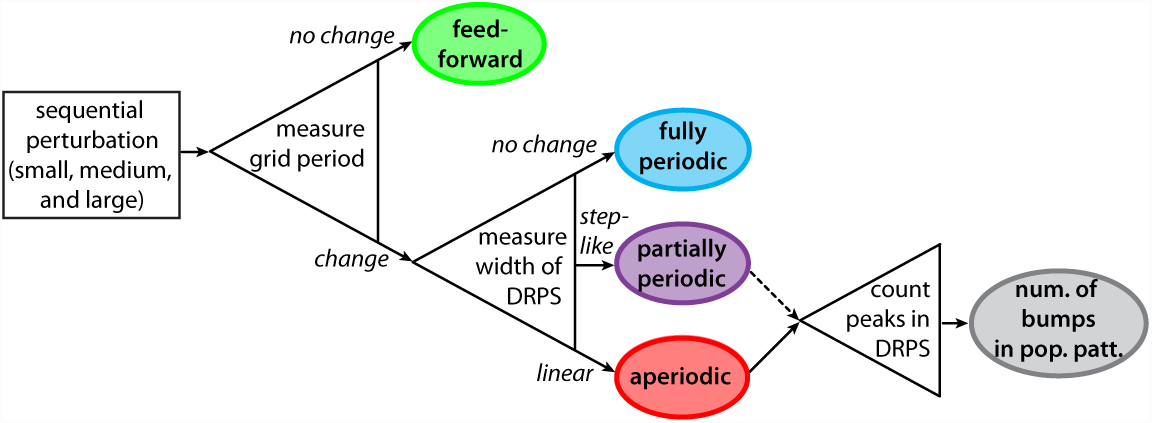
Decision tree for experimentally discriminating circuit mechanisms. For each of three circuit perturbations of increasing strength, both spatial tuning period and relative phase shifts are measured. Recurrent networks are discriminated from feedforward and feedforward-recurrent networks by the effects of the perturbation on spatial tuning period (first open triangle). Different recurrent networks can be discriminated based on how the DRPS width varies with perturbation strength (second open triangle). The number of bumps in the multi-bump population patterns can be inferred by counting the peaks in the DRPS (third open triangle), though, for the partially periodic, only a lower bound on the number of bumps can be established (dotted line).

The demands from experiment are to be able to stably induce a global perturbation in one grid module, and to do so at 2-3 strengths. In all the cases, the term perturbation refers to a small change that leaves the network dynamics qualitatively unchanged while affecting its quantitative properties. The data to be collected are simultaneous recordings from several grid cells as the animal explores a familiar enclosure with no proximal spatial cues over about 20 minutes or more.

First, before applying perturbations, characterize the spatial tuning (periods) of the neurons, as well as cell-cell relationships (the relative spatial tuning phase). Next, apply a series of 2-3 global perturbations of increasing strength. At each perturbation strength, characterize the spatial tuning of cells and cell-cell relationships. A change in the amplitude of the cells’ response across the different perturbations signals that the perturbation is having an effect.

If further there is no change in the spatial tuning period, it follows that the perturbations produced no change in the population pattern and velocity responsiveness, thus the network must be feedforward, Figure 5 (green). Verify that cell-cell relationships remain unchanged across perturbations, as predicted for feedforward networks.

If there is a change in the spatial tuning period, characterize the cell-cell relationships in each perturbation condition. Plot the DRPS from each perturbed condition relative to the pre-perturbation condition, and obtain its width. If the DRPS width increases steadily and linearly with perturbation strength, that implies an aperiodic recurrent architecture, Figure 5 (red). If the DRPS width exhibits a step change, it is consistent with a partially periodic recurrent network, Figure 5 (purple). A DRPS that remains narrowly peaked around zero, with no change in width with perturbation strength, is consistent with a fully periodic network, Figure 5 (blue).

Finally, if the network is either aperiodic or partially periodic, the underlying population pattern has multiple bumps. The number of peaks in the DRPS for each dimension of relative phase bounds from below the quantity 2*M*, where *M* is the number of bumps in the population pattern along that dimension. When the stretch factor *α* times the number of bumps is smaller than 1/2, and if the DRPS is quantal, the number of DRPS peaks equals twice the number of population activity bumps along the corresponding dimension.

## Discussion

### Assumptions

The predictions made here assume that the network activity pattern is stable against rotations. Rotations of the population pattern would induce large changes in the DRPS, obscuring the predicted effects of pattern period expansion in any recurrent network. The fully periodic network is not subject to rotations, but partially periodic and aperiodic networks may be. In experimental data, the cellcell phase relationships between grid cells are indeed very stable across time and environments (Yoon et al., 2013), suggesting that the population activity undergoes no rotation. It is unclear what features of the circuit stabilize the population pattern against rotation; it is possible that slight directional anisotropies in the outgoing connectivity of neurons pin its orientation.

The simplifying observation, that spatial responses may be used to estimate the DRPS, depends on other inputs not being able to overrule the new post-perturbation cell-cell relationships. For instance, external sensory inputs or hippocampal place cells that become associated with particular configurations of grid cells may keep resetting the grid networks to express old relative phase relationships. To avoid this possibility, it may be important to assess post-perturbation cell-cell relationships only in novel environments, for which there are no previously learned associations between external cues, place cell responses, and grid cell activity.

Finally, it is important to note that if in feedforward models one were to include feedback from the grid cell layer back to the spatially tuned inputs (as in Bush and Burgess (2014)), the network would effectively become a type of recurrent circuit, and perturbing the grid cell layer may result in changes in grid period and cell-cell relationships.

### Prior probabilities of different grid cell models being correct

From theoretical arguments, we believe the candidate grid cell mechanisms are not equally probable. In particular, the partially periodic model is difficult to justify from the viewpoint of grid cell development. In Widloski and Fiete (2014), we see that activity-dependent rules acting on spatially informative feedforward inputs can lead to the formation of a network capable of path integration and with grid cell-like tuning. The network, post-development, has aperiodic structure. Under certain conditions, if network weights continue to undergo plasticity after the network has matured enough to expresses recurrent patterning, the network can become fully periodic as neurons with the same spatial phase become wired together (Figure S8). In fact, the addition of relatively weak coupling between neurons in nearest-neighbor activity bumps is sufficient to convert an aperiodic network into what is, functionally if not topologically, a fully periodic network (Figure S8).

Thus, it is possible to imagine mechanisms for the development of the fully periodic and fully aperiodic networks. By contrast, a partially periodic network involves local connectivity which does not depend on a neuron’s spatial phase, but at the same time requires some mechanism for neurons at one end of the network to link with those at the opposite end in way that depends on spatial phase, Figure S1. It is more difficult to imagine a plausible mechanism that can satisfy both constraints. By the same argument, in feedforward models, one would expect the 1D patterned inputs to grid cells to involve fully periodic or fully aperiodic 1D networks.

### Circuit inference through perturbation and sparse activity records: outlook and alternatives

It is interesting to compare the potential of our suggested approach with that of single synapse-level circuit reconstruction (a connectomics approach). A high-quality full-circuit connectome (with signed connections) can specify the topology of the network structure. In other words, it should be possible to reveal whether the circuit is intrinsically “local” (as in the aperiodic network of Figure 1A) (Widloski and Fiete, 2014), partially periodic (with local center-surround-like connectivity and periodic boundary conditions as in Figure 1B), or fully periodic (with center-surround-like connectivity of a width that spans the entire network together with periodic boundary conditions). It may even be possible to infer the locality of structure in the aperiodic network from an unsigned connectome.

Network topology is, however, one ingredient in circuit mechanism: Determining whether the signed connections lead to activity patterning still requires a large amount of inference (for instance, converting the connections into weights and inserting the matrix into an appropriate dynamical model). Even with further inference steps, whether the network actually performs certain functions like velocity-to-position integration or only inherits them is not answerable based on connectomics data. For instance, a network with lateral interactions may generate position-dependent responses *de novo* through integration (Figure 1A-C), or may act only to further pattern inputs that are already spatially tuned (Figure 1D-E) (Kropff and Treves, 2008; Mhatre et al., 2012; Bush and Burgess, 2014). Despite these functional differences, both types of networks have similar connectivity and topologies.

Single neuron-resolution records of activity within a grid cell module can be fruitfully used to understand the dimensionality and relationships of neural responses, but without perturbation, inferring actual connectivity and thus mechanisms from activity is problematic (Roudi et al., 2009; Honey et al., 2009). Hence, activity records do not distinguish between different recurrent models. In short, while connectomics and large-scale recording can provide troves of useful information, they are not sufficient for discriminating between models; as we have shown here, they may also not be immediately necessary.

As we have seen, with a perturbation approach it is possible to localize where integration occurs: if the perturbed area is performing integration, the spatial tuning period is predicted to change. Generally speaking, perturbation modulates the effect of connectivity on dynamics, and the proposed readout is neural activity. This closed-loop approach allows for detailed tests of mechanistic neural models, whose very goal is to relate architecture and dynamics, in a way not easily rivaled by nonperturbative probes of connectivity or activity.

Cooling and other perturbation experiments have been performed in V1 (Michalski et al., 1993; Ferster and Miller, 2000), but they were not as revealing about underlying mechanism as might be possible in grid cells. The reason is twofold: Recurrent models of orientation tuning in V1 are ring models, which are periodic single-bump networks, thus the predicted DRPS after cooling is essentially the same as the prediction for a feedforward network. Moreover, because the orientation response does not arise from integration of a velocity input, the spatial tuning width after cooling is also not predicted to change in a substantial way for recurrent networks. These factors make it harder to discriminate recurrent from feedforward mechanisms from perturbation. The multi-bump tuning of grid cells offers a unique opportunity to use the types of perturbative approaches used in V1 (Michalski et al., 1993; Ferster and Miller, 2000), to obtain unprecedented detail on the local circuit mechanisms that support the complex tuning of cortical cells.

## Methods

### Definitions: Population phase and relative spatial phase

Roman subscripts (e.g., *i* and *j*) refer to individual cells. If cells are arranged topographically based on connectivity, then *i* refers to the location (in neuron body-length units) within the population pattern of the *i*th cell. If the period of the population pattern is *λ*_*pop*_ (again in neuron body-length units), *pop* then the population pattern phase of cell *i*th is 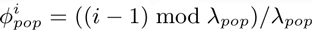 (with the arbitrary choice, made without loss of generality, that neuron 1 has phase 0).

Next, consider the spatial tuning curves of cells *i, j*. Without respect to arrangement in the cortical sheet, let *d*^*ij*^ represent the offset, in meters, of the peak closest to the origin in the cross-correlation of the two spatial tuning curves, and let *λ* be the spatial tuning period (in meters) of the two cells. The relative spatial phase is defined as *δ*^*ij*^ = (*d*^*ij*^ mod *λ*)*/λ*. Phase magnitudes are based on the usual Lee metric, *|δ|* = min(*|δ|,* 1 *- |δ|*). In 2D, the transformation of 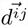 into 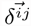 is identical to that described in Yoon et al. (2013) and replicated here in SI. Analogously, using the same procedure, the 2D coordinate of the *i*th cell in the cortical sheet can be transformed into 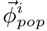 the population phase vector. As noted in Results, *δ*^*ij*^ is easily experimentally accessible; 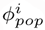, much less so.

### Generation of Figures

Figure 1 is schematic. Figure 2 is generated from ideal (imposed) periodic patterns but without dynamical neural network simulations. In Figures 2, 4A,B, S3, S4, and S7 relative spatial phase is computed for convenience (to save the computational cost of generating spatial tuning curves, then deriving relative phases) from the population phases (thus, by setting 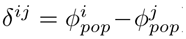). Figures 3, S2, S6, which distinguish between different recurrent architectures, are based on dynamical neural network simulations using the mature grid cell network described in SI. Briefly, the model is a network of excitatory and inhibitory neurons (except in S8 – see figure caption for details), with linear-nonlinear Poisson (LNP) spiking dynamics (Burak and Fiete, 2009; Widloski and Fiete, 2014). For Figure S5, we use Hodgkin-Huxley dynamics. Structured lateral interactions between neurons lead to pattern formation in the neural population. Relative spatial phases are explicitly computed from spatial tuning curves of cells, which are obtained from spike responses to 2-minute long simulated quasirandom trajectories. Velocity inputs drive shifts of the population pattern, resulting in spatially periodic tuning. Only cells from the simulation with good spatial tuning are included in the analysis of relative phase shifts: for fully and partially periodic networks, this means all cells in the network, while for aperiodic networks this means cells in the central 3*/*4 of the network. Since the inhibitory and excitatory populations share similar population patterning and spatial tuning in these simulations, we made the arbitrary choice to display the inhibitory population.

## Supplemental Data

**Figure S1:**
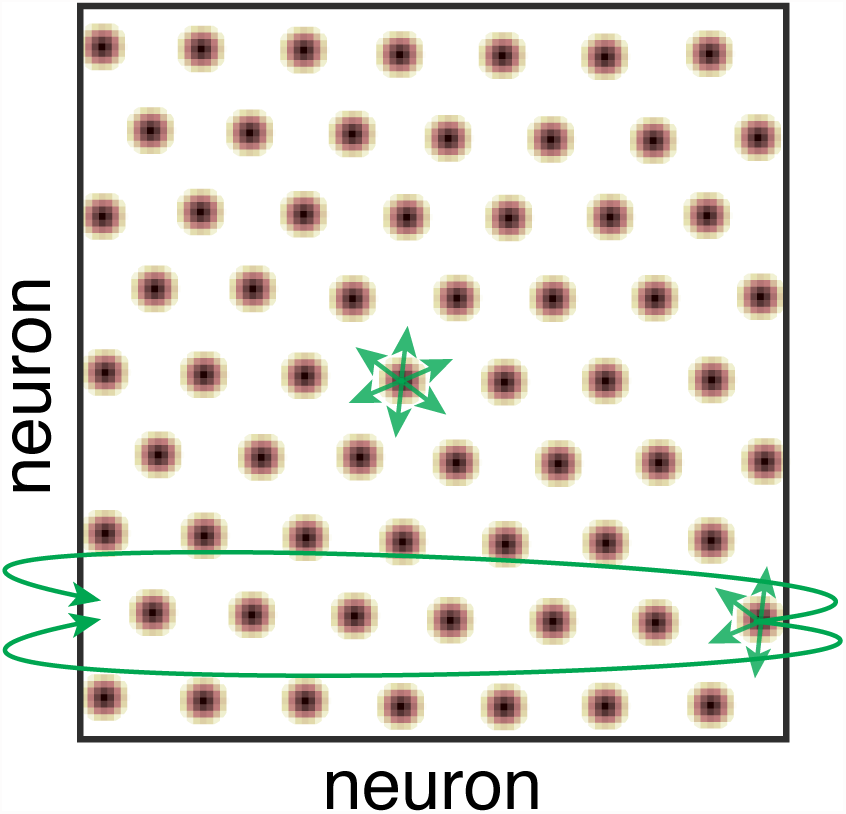
The *a priori* theoretical implausibility of partially periodic networks. Population activity in the cortical sheet (yellow-black blobs), with schematic of connectivity (green). Note that in the bulk of the sheet, connectivity is local and not determined by the periodic activity in the sheet. However, the imposition of periodic boundary conditions requires that some neurons connect with others on the far edge of the sheet. Even if neurons are not topographically organized, the connectivity requires that a planar cortical sheet is somehow intrinsically connected as a torus. Activity-dependent weight changes that are based on the expression of periodic activity patterns could produce a torus-like connectivity, but then if the sheet is not topographically ordered it is likely that neurons in various bumps will connect to each other, producing a fully periodic rather than partially periodic network (see also Figure S8).

**Figure S2:**
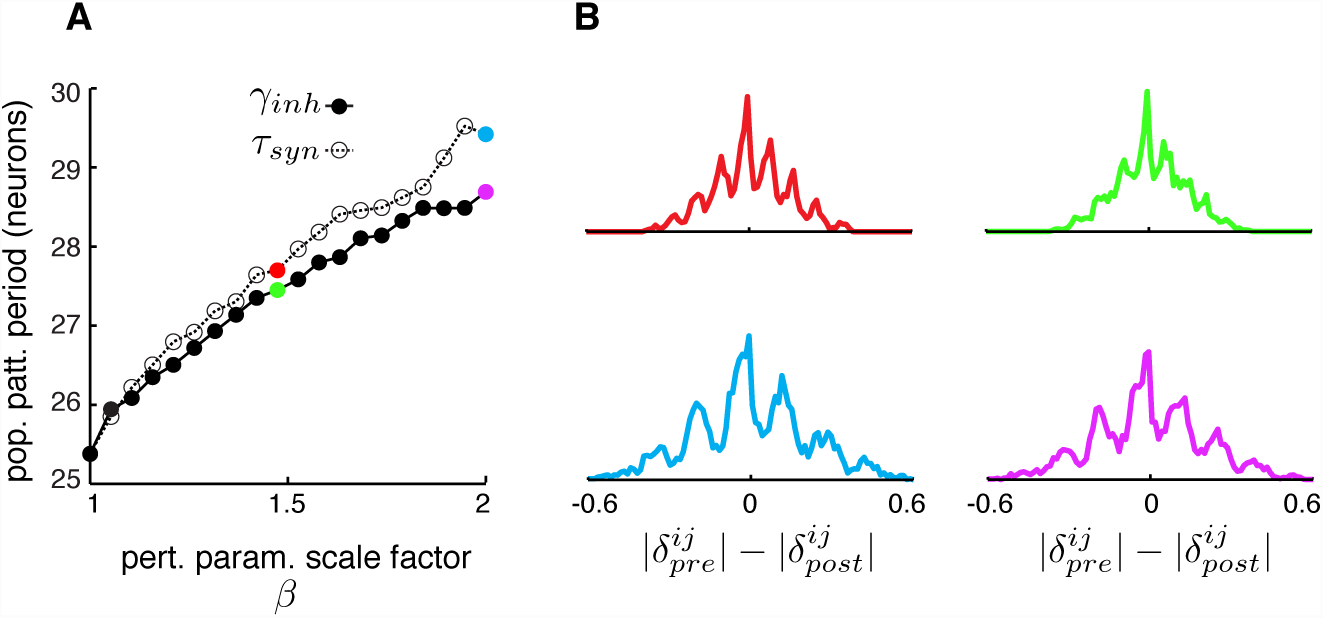
Dynamical simulations of the aperiodic network with LNP dynamics: gradual change in population period and widening of the DRPS with perturbation strength. (A)Change in population pattern period as the inhibition strength (filled circles) or the time-constant (open circles) are scaled up by the factor *β* (inhibition strength 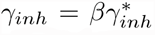 with 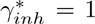; timeconstant 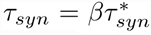, with 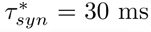) in the 1D aperiodic grid cell neural network (see Experimental Procedures for simulation details and Supplementary Experimental Procedures for definition of population period). (B) DRPS for the data in (A) with respect to the unperturbed network, *β* = 1,, with the size of the perturbations indicated by the corresponding colored dots in (A), with *τ*_*syn*_ (left column) and *γ*_*inh*_ (right column) as the perturbation parameters.

**Figure S3:**
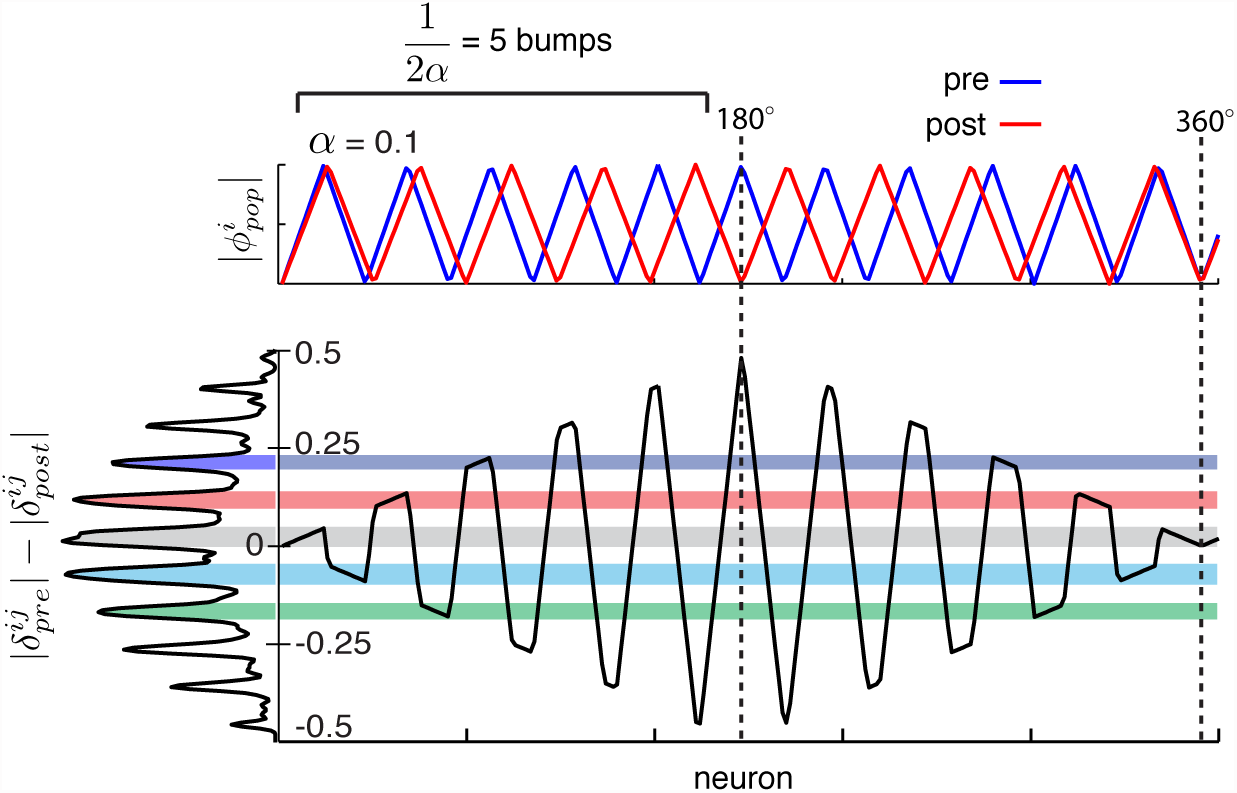
When the 2:1 relationship between number of peaks in the DRPS and the number of bumps in the population pattern breaks down. Top: Schematic of the phase in a population pattern, pre- (blue) and post- (red) perturbation, for a large 1D network with many bumps. If the post-perturbation pattern is aligned to the first bump of the original pattern, the *M* th bump is shifted by an amount *λ*_*pop*_*αM* away from the corresponding bump in the original pattern. When this shift equals *λ*_*pop*_/2, i.e. the perturbed bump is maximally out of phase with the original pattern, there can be no additional (farther out) quantal peaks in the DRPS. Thus, the number of DRPS peaks equals twice the number of bumps in the pattern only when *λ*_*pop*_*αM < λ*_*pop*_/2, or equivalently, when *M α <* 1*/*2. Bottom: Black curve: Difference in the preand post-perturbation phases of cells. At left, the DRPS is aligned vertically with the y-axis of the phase shift plot, so that the origin of the DRPS peaks is more readily apparent. It is clear that once the two patterns reach counter-phase, the locations of DRPS peaks simply repeat. Thus, *M*^*\**^ = 1*/*(2*α*) bumps can accurately be discriminated (per linear dimension of the pattern) for a given stretch factor *α*; when *M > M* ^*\**^, the inference process suffers from systematic underestimation.

**Figure S4:**
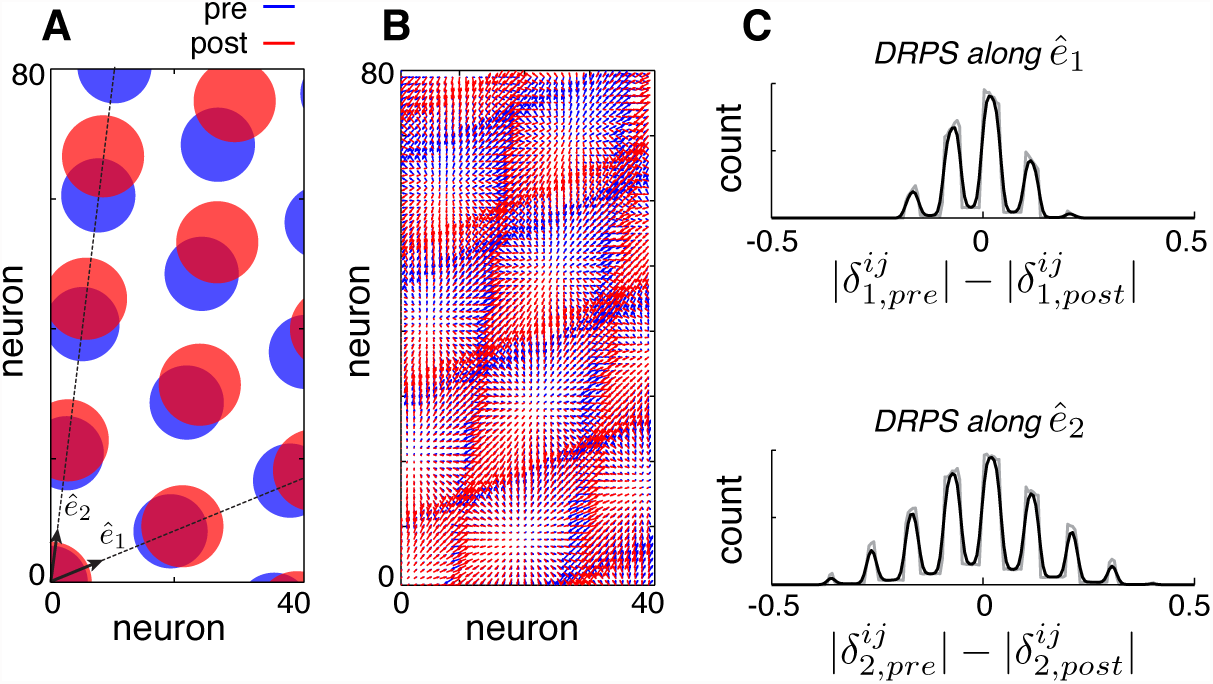
The DRPS in two dimensions. (A) Schematic of 2D population activity pre(blue) and post(red) perturbation (not a dynamical neural network simulation). For illustration, the pattern is depicted topographically and pattern expansion is from bottom left. Subsequent predictions are independent of both choices. Dotted lines: The two principal axes of the pattern. (B) Population phase of each cell, depicted as an arrow (2D phase is a vector). (C) Two relative phase shift histograms computed separately for the two components of the vector phase, along the two principal axes of the lattice (gray: raw data; black: smoothed with 2-bin Gaussian). The DRPS for each phase component resembles the 1D DRPS. Data for each histogram: *n* = (3200 choose 2); bins = 200.*Parameters*: 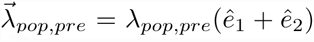, *λ*_*pop,pre*_ = 20 neurons, 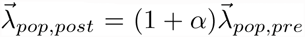, *α* = 0.1, 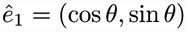, *θ* = 23^*°*^,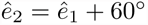, network size: 80 *×* 40 neurons.

**Figure S5:**
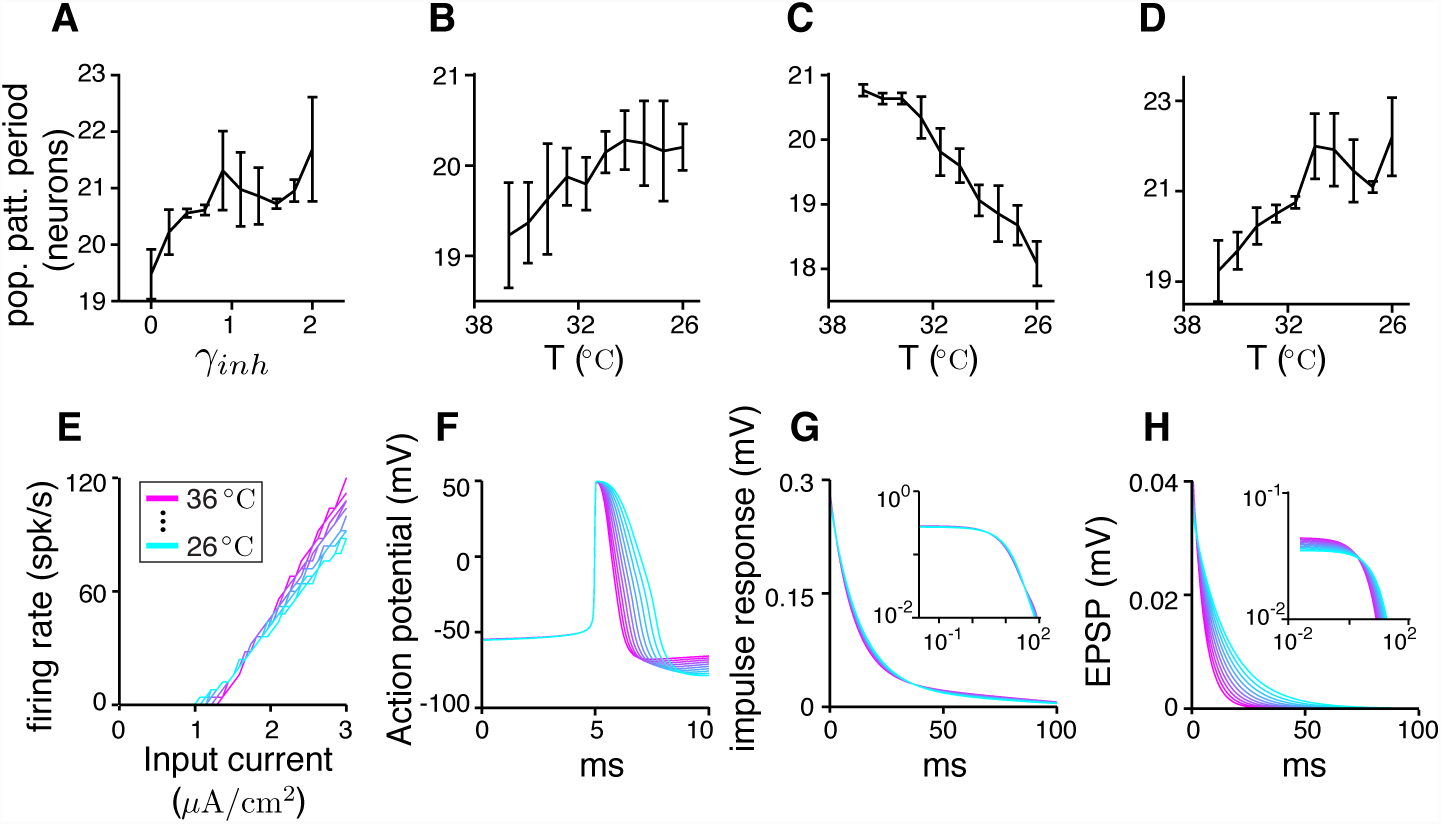
Dynamical simulations of the aperiodic network with HH dynamics: gradual change in population period. (A-D) Population period as a function of perturbation strength (see Supplementary Experimental Procedures for description of HH dynamics and perturbation details). (A) Population period increases gradually with strength of inhibition. Each point is the population period averaged over 10 trials (with error bars equal to the standard deviation). Each trial is 1 second long, in which for the first third of the trial the population pattern is flowed with velocity input equal to 0.3 m/s, and for the remaining time allowed to relax with no velocity input (see Supplemental Experimental Procedures for definition of population period). (B) Population period increases as the temperature is stepped down from 36^*°*^C to 26^*°*^C. (C) Temperature perturbation of only the ionic conductances leads to decreasing population period with decreasing temperature. (D) Temperature perturbation of only the synaptic conductances leads to increasing population period with decreasing temperature. The net result of (C-D) is that the population period increases with decreasing temperature (B). (E-G) Temperature dependencies of single-cell properties, color-coded as a function of temperature. (E) Firing rate as a function of input current, (F) action potential shape, and (G) impulse response (i.e., subthreshold response of membrane potential to current pulse), with log-log plot in inset. (H) EPSP shape as a function of temperature. Though there is a slight decrease in amplitude of the EPSP with temperature (as shown by the log-log plot of the same data in the inset), it is small compared to the effect on the EPSP time constant.

**Figure S6:**
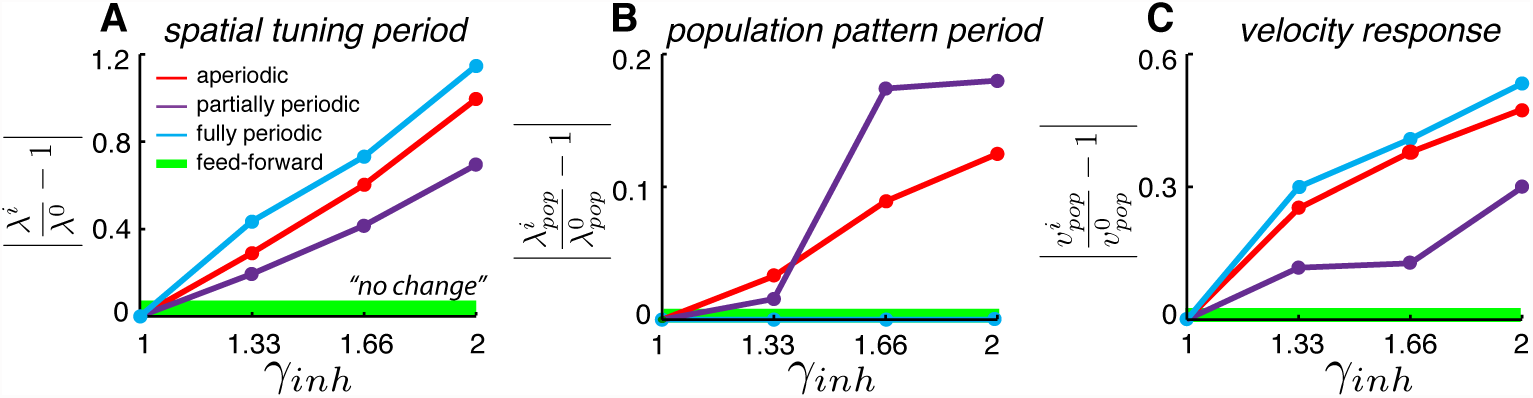
Changes in spatial tuning period in dynamical neural network simulations are due to changes in both the population period and the velocity response of the network. (A) Spatial tuning periods (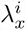 for different perturbation strengths indexed by *i*) of cells in the different recurrent networks, as the strength of inhibition is varied (data as in 3D). (B) The underlying period (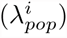) of the population patterns in the same networks for the corresponding strengths of inhibition. (C) The velocity response (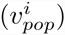) of the networks, or efficacy with which a unit input velocity shifts the phase of the population pattern, as a function of inhibition strength. See Experimental Procedures for simulation details, and Supplemental Experimental Procedures for definition of scores. It is clear that the spatial tuning period (A) is more strongly influenced by the velocity response (C) than by the population period (B).

**Figure S7:**
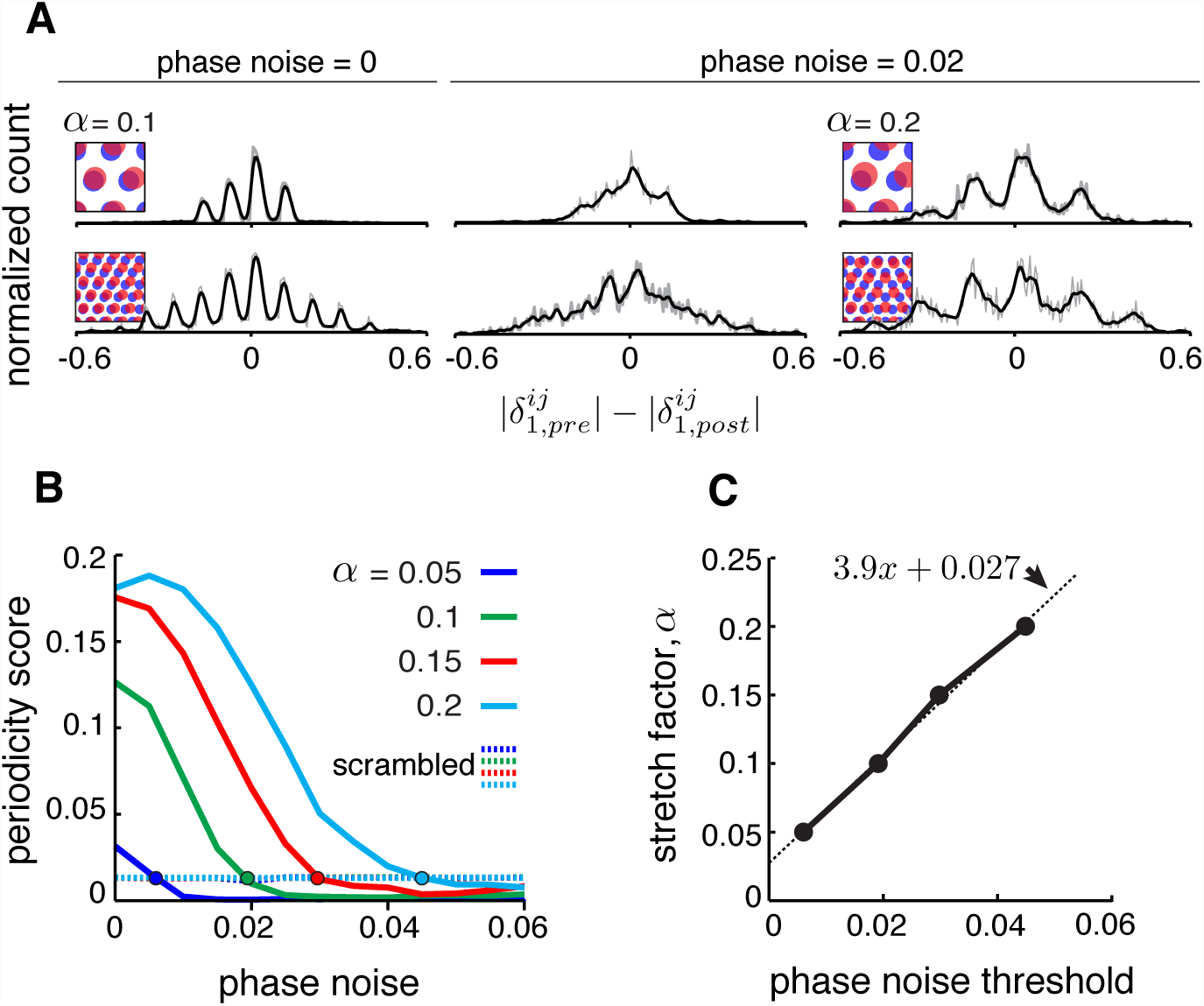
Effects of uncertainty in phase estimation. (A) Copied from Figure 4B. First and second columns: DRPS (200 bins; gray line: raw; black line: smoothed with 2-bin Gaussian) for different numbers of population pattern bumps along the first principal axis of the pattern and for *phase* different amounts of phase noise (noise is sampled i.i.d. from a gaussian distribution, 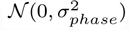, and added to each component of the relative phase vector, 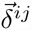; “phase noise” is the same as *σ*_*phase*_). Third column: Same as the second column, except for a larger stretch factor, *α* = 0.2. Note that the peak-to-peak separation has increased so that the individual peaks are discernible. However, for the 5 bump network in the second row, inferring the number of bumps in the underlying population pattern would lead to an underestimate, since *M × α* = 5 × 0.2 *>* 1*/*2. (B) Solid lines: Periodicity score (a measure of how well separated and equidistant are the peaks in the DRPS, and ranges between 0 and 1; see Supplemental Experimental Procedures) as a function of phase noise for 2-bump network in (A), for different values of the stretch factor, *α* (solid lines). Periodicity is measured for the DRPS along the first principal axis. Dashed lines: Same as solid lines, except computed by randomly shuffling the phase vectors post-perturbation. (C) Stretch factor, *α*, as a function of threshold phase noise (defined as the phase noise where the DRPS is indistinguishable from the DRPS when the phase vectors in the post-perturbation condition are reassigned randomly, i.e., the value of the phase noise when the colored curves in (B) cross the respective colored dashed lines).

**Figure S8:**
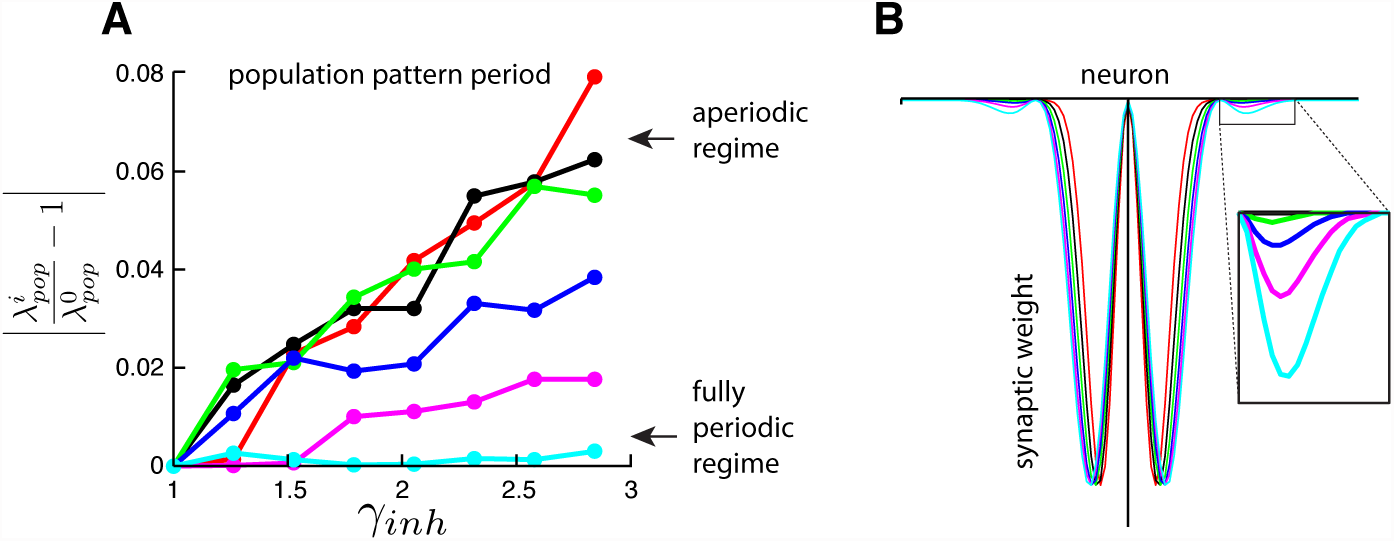
Weakly coupling neurons in different activity bumps in an aperiodic network results in behavior identical to single-bump fully periodic networks. (A) The population pattern in the aperiodic network network exhibits a regime of continuous stretching (black curve) with increasing inhibition strength (*γ*_*inh*_). The ordinate axis is the stretch-factor *alpha*, which quantifies the deviation of the period post-perturbation from that pre-perturbation, normalized by the pre-perturbation period (See Supplemental Experimental Procedures for definition of population period). However, adding even very weak synaptic connections between neurons in adjacent activity bumps in the aperiodic network (B) transforms the network into one that will not stretch at all (cyan curve), like the single-bump fully periodic network. The two constructions (the single bump network and the aperiodic network with the addition of between-bump connections) would be mathematically the same if there were strong coupling between all neurons of the same activity phase in the network; this numerical result shows that the addition of very small weights that reflect the periodicity of the population activity (but are nevertheless largely local within the network in the sense that only neighboring bump neurons are connected, not neurons in remote bumps) already transforms the aperiodic network into an effectively fully periodic, single-bump network. *Simulation details*: The network connectivity is as a hybrid of the aperiodic network in Burak and Fiete (2009) with the fully periodic network of Fuhs and Touretzky (2006) (note that, while the model of Fuhs and Touretzky (2006) does not have explicit periodic boundary conditions, the multimodality of the synaptic weights couples adjacent activity bumps so that the network acts as a single-bump, fully periodic network). The dynamics are LNP-based (see Supplementary Experimental Procedures) and driven with inputs simulating animal motion at constant speed (v = 0.3 m/s) for 10 seconds. There are only two populations (call them R and L), distinct in their directional preferences 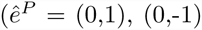 for the R and L populations, respectively) and output synaptic asymmetries (see below). The shifted output weight profiles are sinusoids with gaussian envelopes, the latter which constrain the non-locality of the projections. For a narrow gaussian envelope, the weights resemble the purely local, center-surround profiles of Burak and Fiete (2009), whereas for wide gaussian envelopes, the weights resemble the non-local, multimodal projections of Fuhs and Touretzky (2006). The weights going from population *P*′ to *P* and from cells i and j, are given by 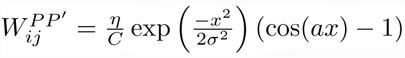, where *x* = *i - j* + Δ (Δ = *±*1 for *P′* = *R*(+) and *P*′ = *L*(+)), *η* is a scaling factor that modulates the amplitude of the weights, 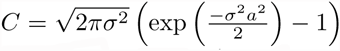 is a normalization factor, *σ* determines the width of the is a normalization factor, *σ* determines the width of the gaussian envelope, and *a* determines the period of the underlying sinusoid. *Parameters*. *N*_*R*_ = *N*_*L*_ = 200 neurons; CV = 0.5; *dt* = 0.5 ms; *τ*_*syn*_ = 30 ms; *g*^0^ = 50; *g*^0′^ = 0; *β*^*vel*^ = 1; 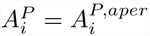; *a* = 2*Π/*20; *η* = 200; *σ* = 4*→*12;

## Supplemental Experimental Procedures

### Neural network simulations

Below, we describe the two different neural dynamics models used in the paper: the linear-nonlinear-Poisson (LNP) model and the Hodgkin-Huxley conductance model.

Roman subscripts (e.g. *i, j*) refer to individual cells within population *P*. The population index *P* can take the values {I, E^*R*^, E^*L*^}. Integration in all simulations is by the Euler method with time-step *dt*.

#### Linear-Nonlinear-Poisson dynamics (all figures except Figure S5)

The LNP model we use is identical to that used in Widloski and Fiete (2014). Given a time-dependent summed input 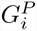 to the (*P, i*)th cell, the instantaneous firing rate of the cell is

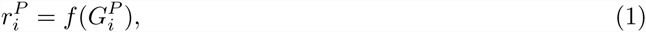

with the neural transfer function *f* given by

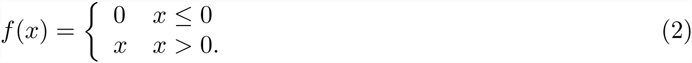

Based on this time-varying firing rate, neurons fire spikes according to an inhomogeneous (sub-Poisson) point process with a coefficient of variance of CV = 0.5 (see Burak and Fiete (2009) and Widloski and Fiete (2014) for details on generating a sub-Poisson point process).

The time-dependent activation 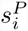 of synapses from the (*P, i*)th cell is given by

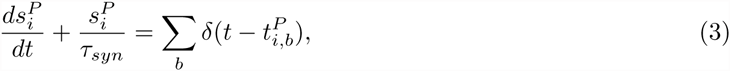

where 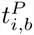 specifies the time of the *b*th spike of the cell and the sum is over all spikes of the cell.

The total input 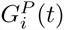 into the (*P, i*)th cell is given by

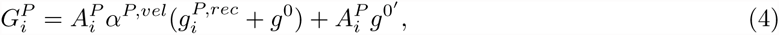

where *g*^0^ (*g*^0^=50 for the E and I populations) and *g*^0′^ (*g*^0′^ =15 for the E population; *g*^0′^ =0 for the I population) are small, positive, constant bias terms common to all cells, 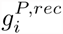 are the recurrent inputs, *α*^*P,vel*^ are the velocity inputs, and 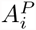 is an envelope that either suppresses activity near the network boundaries for the *aperiodic* network, or is flat and equal to unity for the *periodic* networks (see below). The recurrent input is

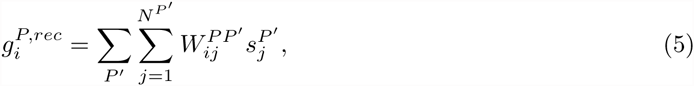

where 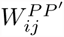are the recurrent weights and *δ* is the Kronecker delta function. The form of the envelope function, 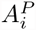, depends on the boundary conditions of the network. For *aperiodic* networks, the envelope shape is a 1D version as that given by Burak and Fiete (2009):

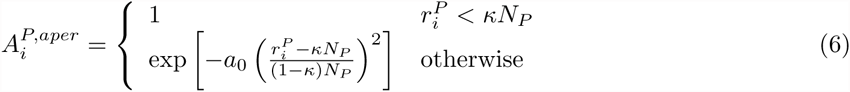

where *N*_*P*_ is the size of the network, 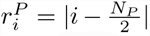, *κ* = 0.3 determines the range over which tapering occurs, and *a*_0_ = 30 controls the steepness of the tapering. For *periodic* networks, the envelope is flat:

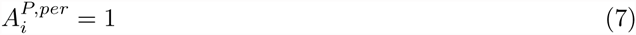

All cells in the *P*th population (with preferred direction given by the unit vector 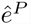) receive a common velocity input:

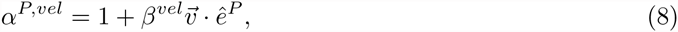

where 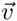 is instantaneous velocity of the animal and *β*^*vel*^ sets the gain of the velocity input; 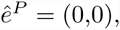 (0,1), (0,-1) for the I, E^*R*^, E^*L*^ populations, respectively. The velocity input, unless otherwise noted, is based on a 2-minute quasi-random trajectory derived with an algorithm identical to that described in Widloski and Fiete (2014). Over the course of these trajectories, the stochastic dynamics leads to drift in the path-integrated estimate of animal location if uncorrected. To minimize this drift, the pattern phase is reset whenever the animal is in the vicinity of one of the 5 ‘landmarks’ evenly spaced throughout the environment. During each encounter with a landmark, the pattern phase is corrected via strong feedforward inputs that impose a snapshot of the pattern at its “correct” phase; “correct” pattern snapshots are captured from the population pattern during the animal’s initial encounter with each of the landmarks.

*Temperature/neuromodulation of LNP dynamics*. Temperature-dependent modulations are modelled as a simple rescaling of the synaptic activation time constant, *τ*_*syn*_. Modulations of network inhibition are modelled as a gain change in the efficacy of the synaptic weights projecting from inhibitory neurons, i.e., *W* ^*P*^ ^*I*^ *← γ*_*inh*_*W* ^*P*^ ^*I*^, where *γ*_*inh*_ is the strength of inhibition.

#### Hodgkin-Huxley dynamics (only used in Figure S5)

The model we use is identical to the reduced Hodgkin-Huxley “regular spiking (RS)” model of cortical neurons, as described in Pospischil et al. (2008), supplemented with synaptic dynamics. The dynamics of the membrane potential of the (*P, i*)th neuron is given as

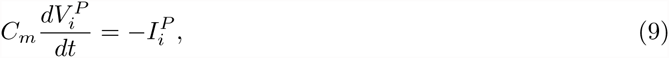

where 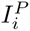 is the summed input current and *C*_*m*_ is the capacitance of the membrane. The summed input current is given as

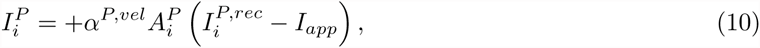

where the first term represents currents related to the ionic membrane conductances and the second and third terms represents synaptic and external conductances, respectively, gated by velocity inputs, *α*^*P,vel*^, and an envelope function, 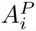. The ionic current has the following form:

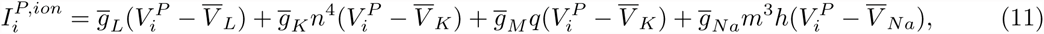

where the 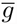’s are the maximum conductance values and the 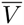’s the reversal potentials of the leak conductance (L), fast (K) and slow (M) potassium conductances, and the sodium conductance (Na). The dynamics and parameter settings of the gating variables *n, m, q, h* are described in Pospischil et al. (2008) (note that we have replaced the “p” gating variable of Pospischil et al. (2008) with “q”). The synaptic current based on recurrent connections within network is

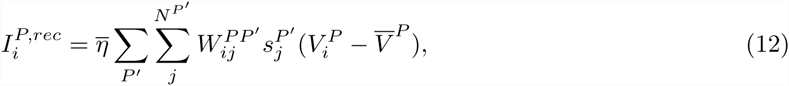

where 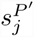 is the synaptic activation of the (*P ^l^, j*) neuron (which has the same dynamics as described above in equation (3) – here, we define the time of a spike elicited by the *j*th neuron, 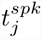, as when the voltage moves from below 0 mV to above it in a single-time step, within the interval (*t, t* + Δ*t*)), 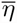 is a synaptic scaling factor shared by all synaptic weights, and 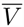 is the synapse-specific reversal potential (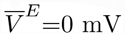 and 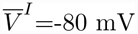).

*Temperature/neuromodulation of HH dynamics*. To simulate temperature-dependent modifications, we used separate *Q*_10_ factors to modulate the time constant 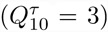 and amplitudes 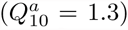 of the ionic/synaptic conductances (Hodgkin et al., 1952; Katz and Miledi, 1965). At temperature T (^*°*^C), the conductance amplitudes 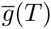 and time constants *τ* (*T*) have the following form:

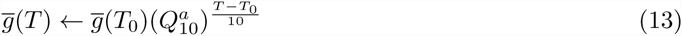

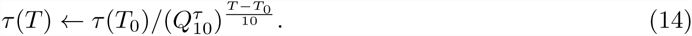

The conductance amplitude modulation was applied specifically to 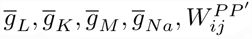. The conductance time constant modulation was applied to the gating variable time constants *τ*_*n*_, *τ*_*q*_, *τ*_*m*_, *τ*_*h*_ (for gating variable *x*, the time constant *τ*_*x*_ is defined as *τ*_*x*_ = 1*/*(*α*_*x*_ + *β*_*x*_), where *α*_*x*_ and *β*_*x*_ are the rate constants governing the gating variable’s dynamics – see Pospischil et al. (2008)) and the synaptic time constant *τ*_*syn*_. For temperature perturbations of the ionic conductances only, 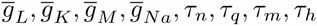 change with temperature, while 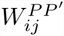 and *τ*_*syn*_ =16 ms are held constant. For temperature perturbations of the synaptic conductances only, 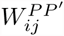 and *τ*_*syn*_ change with temperature, while the ionic conductance properties are held fixed.

The effects of specific neuromodulators targeting the inhibitory synapses was modelled in exactly the same as for the LNP model.

#### Synaptic weights for network of excitatory and inhibitory neurons (all figures except Figure S8)

The detailed synaptic weights used in the simulations are based on the developmentally-inspired hardwired weights with aperiodic boundary conditions described in the SI of Widloski and Fiete (2014), and therefore can be viewed as the plausible culmination of a developmental process. Compared to the LNP-based model used in Burak and Fiete (2009), we chose to implement the model in Widloski and Fiete (2014) because it is more realistic, incorporating separate populations of excitatory and inhibitory cells; however, both models give qualitatively similar results. (Note that while the description of the weights below is different than that specified in Widloski and Fiete (2014) in order to enable flexibility in setting the network boundary conditions, the weights used for the aperiodic network are identical to those specified in Widloski and Fiete (2014).) The weights from population *P*^*l*^ to *P*, between cells i and j, are described as follows:

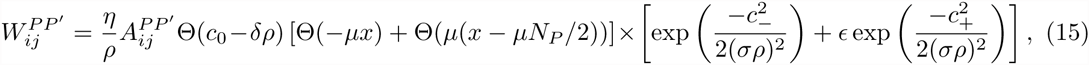

where *x* = *i γj*, 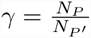 (*N*_*P*_ is the size of population *P*), Θ is the Heaviside function (Θ(*x*) = 0 for *x <* 0 and is 1 otherwise), *c*_0_ = *Ψ*(*x*) and *c*_*±*_ = *Ψ*(*x±*Δ*ρ*) where *Ψ*(*x*) = min (*N*_*p*_ - *|x* mod *N*_*p*_*|, |x* mod *N*_*p*_*|*), and 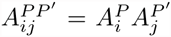, where 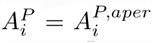 for *aperiodic* networks and 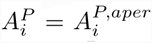 for *periodic* networks (see above for definition of 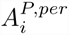 and 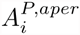). *ρ* is a scale factor that controls the width of the synaptic weights, and therefore the number of bumps expressed in the pattern, whereas *η* is a synaptic scaling factor that modulates only the amplitudes.

(A note on terminology: the *partially periodic* network has an overall topology that resembles the *periodic* network of Burak and Fiete (2009). In our usage in the present work, periodic refers to a fully periodic network, in which the periodicity of connections matches that of activity pattern, whereas in the partially periodic network, the bulk of connectivity does not reflect the periodicity of the population activity pattern.)

#### Simulation parameters

*Aperiodic network with LNP dynamics*.

*P* = *E*_*L*_, *E*_*R*_, *I*; *N*_*E*__*L*_ = *N*_*E*_*R*__ = 400 neurons; *N*_*I*_ = 160 neurons; CV = 0.5; *dt* = 0.5 ms; *τ*_*syn*_ = 30 ms*; *β*^*vel*^ = 2; 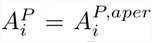; *ρ* = 2.2. *γ*_*inh*_ = 1*;

*E*^*L*^ *→ I* (i.e., *W ^IE^L*): *η* = 21; *E* = 0; Δ = −1; *σ* = 2; *μ* = 0; *δ* = 0;

*E*^*R*^ *→ I*: *η* = 21; *E* = 0; Δ = 1; *σ* = 2; *μ* = 0; *δ* = 0;

*I → E*^*L*^: *η* = 8; *E* = 0; Δ = 4; *σ* = 5; *μ* = −1; *δ* = 3;

*I → E*^*R*^: *η* = 8; *E* = 0; Δ = −4; *σ* = 5; *μ* = 1; *δ* = 3;

*I → I*: *η* = 24; *E* = 1; Δ = 2; *σ* = 3; *μ* = 0; *δ* = 3;

(* indicates that parameters can change through perturbation)

*Partially periodic network with LNP dynamics*.

Same parameters as aperiodic network, except that 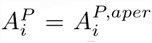, and *ρ* = 2.2.

*Fully periodic network with LNP dynamics*.

Same parameters as aperiodic network, except that 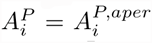, and *ρ* = 22.

*Aperiodic network with HH dynamics*.

All ionic conductance parameters are identical to those described in Pospischil et al. (2008) for the RS model; as noted there, the parameters are set to values corresponding to a temperature of *T*_0_ = 36^*°*^*C*. *N*_*E*__*L*_ = *N*_*E*__*R*_ = 400 neurons; *N*_*I*_ = 160 neurons; *dt* = 0.025 ms; *τ*_*syn*_ = 15 ms*; *β*^*vel*^ = 0.8; *C*_*m*_ = 1 *μ*F/cm^2^; 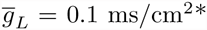; 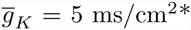; 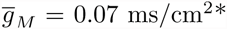; 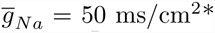; 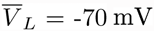; 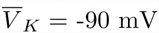; 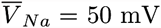; *I*_*app*_ = = 3 *μ*A/cm^2^; 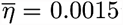; 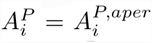; *ρ* = 2.2. *γ*_*inh*_ =*i i* 1*; Synaptic weights are identical to those described for the aperiodic network with LNP dynamics up to a constant, 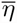. (* indicates that parameters can change through perturbation)

#### Measures used in main text

*Bootstrap resampling and phase uncertainty.* Given an original spike map of *M* total spikes (with locations) from one cell, we created a new spike map of *N* (*N < M*) total spikes, by picking spikes (with their corresponding location coordinates) from the original map one at a time, at random, and with replacement. The same was done for a second, simultaneously recorded cell. From these sampled spike trains for a pair of cells, we estimated relative phase (by computing the location of the peak closest to the origin in the cross-correlation of the spatial maps of the two cells, as in Yoon et al. (2013)). The procedure was performed 100 times, generating 100 bootstrapped relative phase estimates per cell pair. Phase uncertainty was measured as the mean of the magnitudes of the bootstrapped relative phase estimates.

*Spatial tuning curves.* For a given cell and trajectory, we build a histogram of spike counts at each location (bin size = 1 cm), then normalize the count in each bin by the amount of time spent in it. The normalized histogram is smoothed by convolution with a boxcar filter (width = 5 bins) to yield a spatial tuning curve.

*Spatial tuning period.* The spatial tuning period is measured as the inverse of the spatial frequency with the highest peak in the power spectrum of the spatial tuning curve (excluding the peak at 0 frequency).

*Population period.* Given the last 500 snapshots (frames) of the population pattern from a given trial, the population period is measured as followed: For each frame, measure the inverse of the frequency with the highest peak in the power spectrum (as with the spatial tuning period) of the population pattern. The population period is the average of these estimates.

*Velocity response.* Velocity response is measured as the translation speed (neurons/sec) of the network pattern to fixed input velocity, computed by tracking the displacement of the pattern for 10 seconds, smoothing the resulting trajectory with an 4-second moving average filter, and then measuring the average speed of the middle-half of the trajectory.

*Periodicity score for the DRPS.* We smooth the histogram of relative phase shifts (by convolution with a 2-bin Gaussian kernel) and normalize it (by mean subtraction and division by the standard deviation). Next, we compute the power spectrum, rescaling the result by 2*/L*^2^, where *L* is the number of bins in the histogram (*L* = 200). The periodicity score is set to be the power of the largest-amplitude non-zero frequency component in the scaled power spectrum. This score returns 1 if the DRPS is a pure sinusoid. It returns 0 if the DRPS is flat and returns an average value of *<* 0.2 if the DRPS were constructed bin by bin by sampling iid from a uniform distribution on the unit interval.

*2D relative phase.* The displacement vector *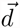 is* converted into a 2D phase 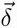 according to 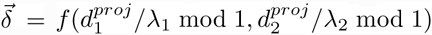, where 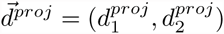 is the oblique projection of 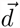 onto the principal vectors 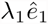and 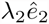, and

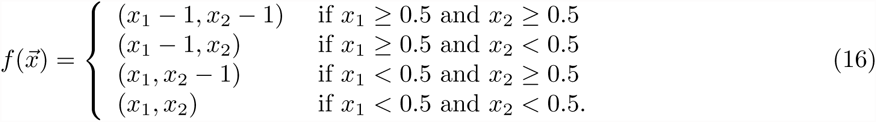

Relative phase magnitude is given by

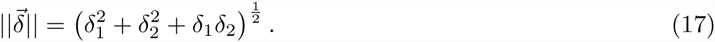

